# Decentralized Distribution-sampled Classification Models with Application to Brain Imaging

**DOI:** 10.1101/576108

**Authors:** Noah Lewis, Harshvardhan Gazula, Sergey M. Plis, Vince D. Calhoun

**Affiliations:** The Mind Research Network, Albuquerque, NM; Department of Computer Science, The University of New Mexico, Albuquerque, NM; Department of Electrical and Computer Engineering, The University of New Mexico, Albuquerque, NM

## Abstract

**background:** In this age of big data, large data stores allow researchers to compose robust models that are accurate and informative. In many cases, the data are stored in separate locations requiring data transfer between local sites, which can cause various practical hurdles, such as privacy concerns or heavy network load. This is especially true for medical imaging data, which can be constrained due to the health insurance portability and accountability act (HIPAA). Medical imaging datasets can also contain many thousands or millions of features, requiring heavy network load.

**New Method:** Our research expands upon current decentralized classification research by implementing a new singleshot method for both neural networks and support vector machines. Our approach is to estimate the statistical distribution of the data at each local site and pass this information to the other local sites where each site resamples from the individual distributions and trains a model on both locally available data and the resampled data.

**Results:** We show applications of our approach to handwritten digit classification as well as to multi-subject classification of brain imaging data collected from patients with schizophrenia and healthy controls. Overall, the results showed comparable classification accuracy to the centralized model with lower network load than multishot methods.

**Comparison with Existing Methods:** Many decentralized classifiers are multishot, requiring heavy network traffic. Our model attempts to alleviate this load while preserving prediction accuracy.

**Conclusions:** We show that our proposed approach performs comparably to a centralized approach while minimizing network traffic compared to multishot methods.

**Highlights:**

- A novel yet simple approach to decentralized classification
- Reduces total network load compared to current multishot algorithms
- Maintains a prediction accuracy comparable to the centralized approach

## 1 Introduction

Due to advances in analytic tools, researchers have the opportunity to assemble informative models given large datasets. Because of this, research institutes and hospitals are increasing collaborative efforts [Ming et al., 2017, Plis et al., 2016, Carter et al., 2015, Thompson et al., 2014]. The simplest and easiest method is for institutes to share data, which typically involves an individual group downloading data from various sources and performing large, centralized analyses. Such an approach is limited in what data can be shared openly due to HIPAA or other regulatory concerns. More specifically, these institutions need to protect the personal information of patients which can be compromised when transferring or centralizing the data. While this can be addressed by anonymizing data, in some cases, notably when there are many local sites, this is not sufficient. Another issue is that large computational resources and extensive download time is typically required in a centralized approach, which is very problematic for neuroimaging datasets, which can contain hundreds or thousands or subjects, with many thousands or millions of features. Decentralized models may solve both of these issues by eliminating the requirement of data transfer.

There is a current body of various decentralized models [Gazula et al., 2018, Saha et al., 2017, Wojtalewicz et al., 2017, Baker et al., 2015], and more specifically, decentralized neural networks [Lewis et al., 2017] and support-vector machines (SVM)[Forero et al., 2010]. However, these models are multishot, meaning they pass statistical information many times during the training process, which can require a great deal of network traffic. The multishot neural network, or decentralized-data neural network (dDNN) [Lewis et al., 2017] requires heavy network traffic at least once every epoch, or one full iteration through the entire dataset during the training process. This is because the dDNN model passes all gradient information from local sites to a centralized location after every epoch, then calculates the average of these gradients, and passes the averaged gradients to the local sites. As neural networks can require many thousands of epochs, the overall network traffic would be unmanageable for neuroimaging data, which can contain hundreds of thousands of features. This same problem occurs for multishot SVMs, which also require a high number of steps in which gradients are passed between local sites.

In this research, we attempt to mitigate these issues for certain classifiers by introducing a singleshot method. Singleshot methods require statistical information to be passed only once, either before or after the local models have been trained. In our case, statistical information is passed to the local sites, and then each site trains separately. The statistical information is an estimated distribution of the local data, which is comprised of the per-feature mean and a covariance matrix of the features. We refer to this model as a decentralized distribution-sampled classifier (dDSC). This use of statistical inference to estimate new samples for decentralized modeling is applied to both neural networks (dDS-NN) and SVMs (dDS-SVM) to show efficacy in use with multiple classification models. We quantify the data at each local site by building local distributions using a Gaussian mixture model (GMM) and pass these distributions to the remaining local sites which will then be used in training models at the local sites. Each local site combines artificial data sampled from the given distributions with locally available data to train the models. We demonstrate the efficacy of dDS-NN and dDS-SVM on two datasets.

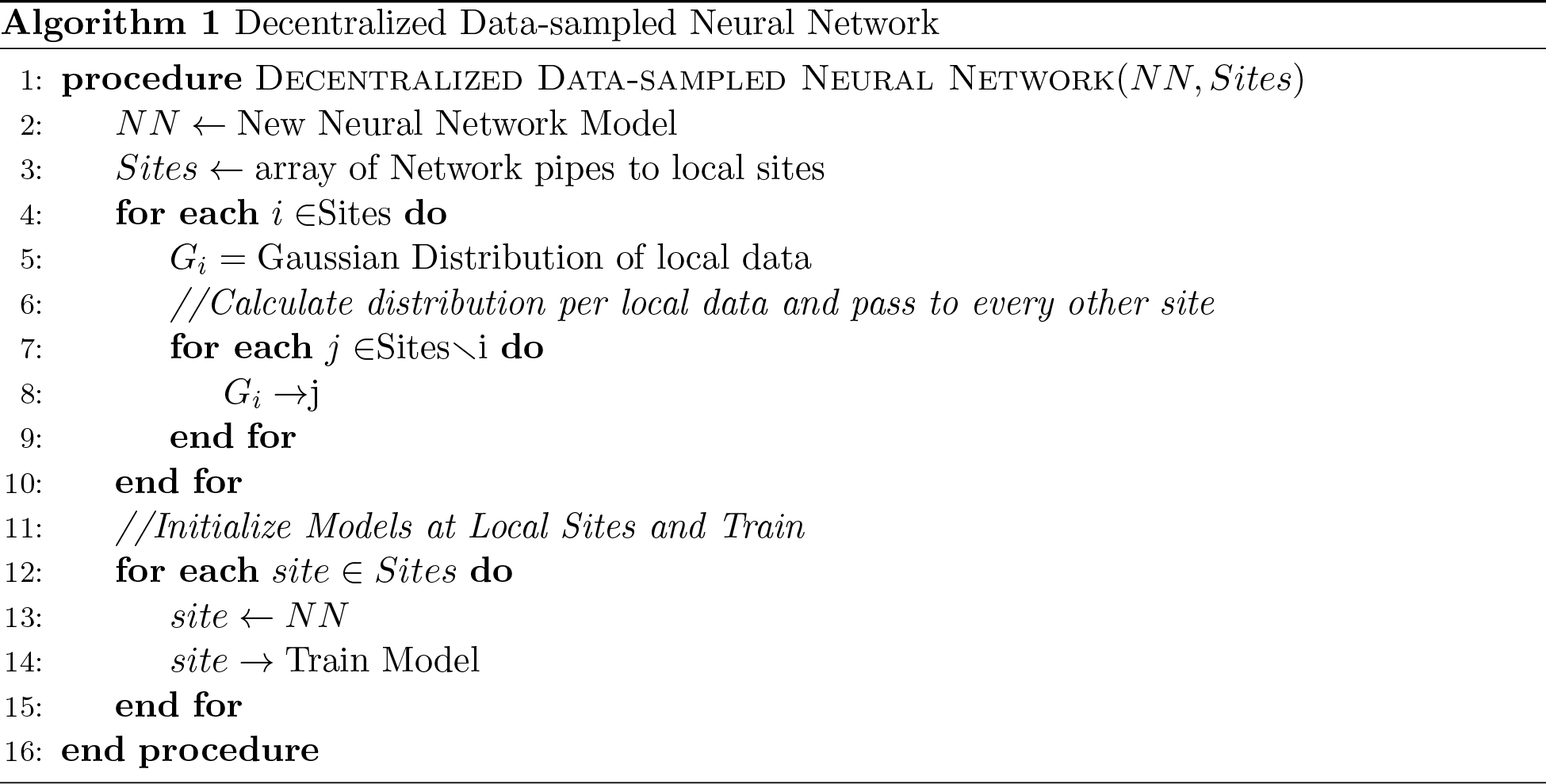

## 2 Methods

### 2.1 Multishot decentralized modalities

In the previous multishot models, dDNN and the consensus-based SVM, high-level statistical information (i.e. gradients) are passed between local sites many times during the training of the models. However, this requires a high traffic load and as the number of training iterations increases, the chance of network failure also increases. The dDNN aggregates the local gradients every iteration (or epoch through the data), averages the gradients, and passes these updated gradients to the local sites. The multishot SVM uses the alternating direction method of multipliers (ADMoM) to accumulate the updating parameters, or the model weights, [Forero et al., 2010]. These model weights are used, as in a non-decentralized model, to update the Lagrangian multipliers. This process is repeated as many times as necessary to complete the training.

### 2.2 Statistical inference models

Our approach for singleshot classifiers-dDSC gathers statistical information about the datasets, rather than the models as is the case in the multishot algorithms, at the local sites and passes this information between the sites before the models are trained. We use a GMM to estimate the distribution of the local site data for each class. Once the distribution is gathered from the model for each site, this distribution is passed to the other sites. The other sites then draw artificial samples from the remaining sites’ distributions and trains their own model on both locally available data as well as the artificial samples. This approach also shows a much smaller amount of network traffic, as the mixture model is transferred once, with a polynomial relationship to the number of input features. This is the case for both the neural network and SVM methods.

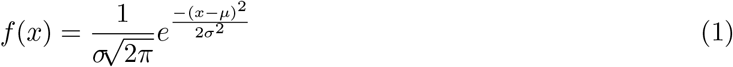

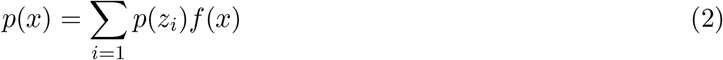

### 2.3 Gaussian Mixture Model

A GMM uses the expectation-maximization (EM) algorithm to fit the data to a distribution. The EM algorithm begins by, for each class, creating a random Gaussian distribution, or randomized mean and variance. Then, the probability that each data point is within a given distribution is calculated using the Gaussian equation (Equation 2). Finally, the mean and variances of the distributions are recalculated based on the probability of each data point being a member of a given distribution. Once the parameters have been re-estimated, the process starts over again, and the data points are reassigned to a distribution given the updated means and variances. This process continues until the model converges to a given maximum likelihood estimation.

GMMs use covariance matrices to store the variances of the features, which determine the contours of the distribution in *N*-Dimensional space. The covariance matrices can be represented in different ways, the most common being a full-covariance matrix; in which the variance of all pairs of features is stored in the matrix. Another representation of covariance matrices is a diagonal matrix, in which the variances of only individual features are stored in memory. This distinction is important for our work as it determines the network traffic of the model. When analyzing the MNIST dataset, we use a full covariance matrix as the feature set is relatively small compared to the sMRI dataset. However, in the case of the sMRI data, we use a diagonal matrix due to the sheer size of the feature space and to show that the dDSC also works with a very small subset of the distribution information, which reduces total network traffic.

### 2.4 Experiments

The experiments use two different datasets: the mixed national institute of standards and technology (MNIST) dataset of handwritten numbers [LeCun et al., 2010], and a set of real-world structural magnetic resonance images (sMRI) of schizophrenia patients and healthy subjects from an aggregated multisite dataset [Potkin et al., 2008, Gollub et al., 2013, Hanlon et al., 2011, Aine et al., 2017]. We tested the models’ performances on MNIST with three cases: the data is uniformly and randomly distributed across three sites, three sites have access to only certain classes, and the datasets are uniformly and randomly distributed across 20 sites. We also tested the models via the sMRI data in two cases: the data is uniformly distributed across four sites at random, and four sites have access to only certain classes. A full break-down of the experiments can be seen in table 1.

**Table 1:**
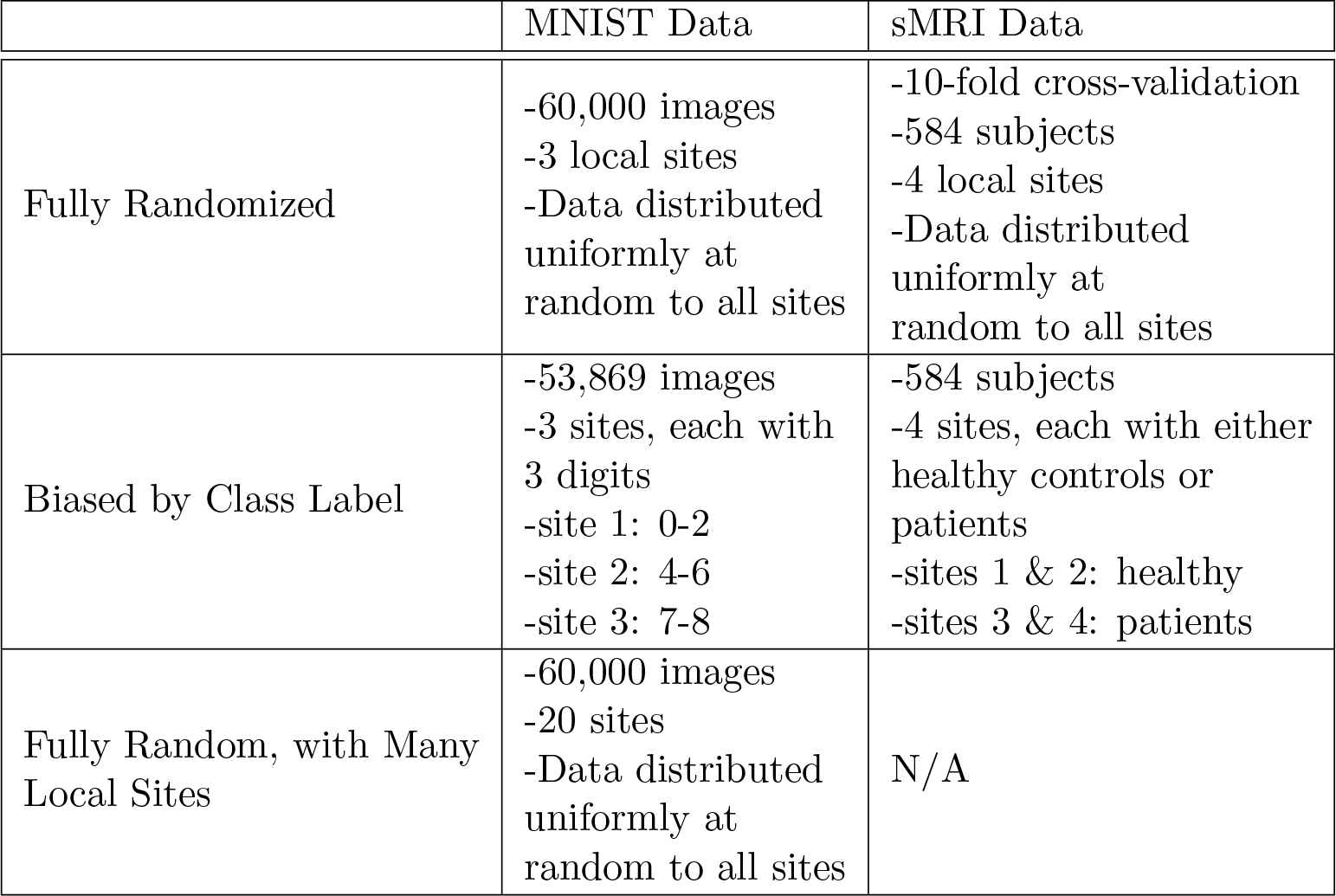
The break-down of the experiments. The rows are the two different datasets, and the columns are the three experiments. Each cell explains the important details of an experiment for a given dataset.

|  | MNIST Data | sMRI Data |
| --- | --- | --- |
| Fully Randomized | -60,000 images<br>-3 local sites<br>-Data distributed uniformly at random to all sites | -10-fold cross-validation<br>-584 subjects<br>-4 local sites<br>-Data distributed uniformly at random to all sites |
| Biased by Class Label | -53,869 images<br>-3 sites, each with 3 digits<br>-site 1: 0-2<br>-site 2: 4-6<br>-site 3: 7-8 | -584 subjects<br>-4 sites, each with either healthy controls or patients<br>-sites 1 & 2: healthy<br>-sites 3 & 4: patients |
| Fully Random, with Many Local Sites | -60,000 images<br>-20 sites<br>-Data distributed uniformly at random to all sites | N/A |

### 2.5 MNIST Experiments

For both MNIST experiments, the images are vectorized from 28×28 pixel images into vectors of size 784. The labels are one-hot-encoded into a vector of size 10 for each image. For all experiments, we tested the models using the test set established by the dataset creators.

In the first dDS-NN experiment, we randomly select 20,000 images from the entire dataset of 60,000 images for each of the three sites. This process is used for the dDNN and dDS-NN model.

However, the centralized model have one site with access to all 60,000 images. All of the neural networks use the same architecture: two hidden layers of size 256, a learning rate of 0.001, and batch normalization is also used. The models are initialized with the Xavier initialization algorithm (Glorot, 2010). The models are run for 1000 epochs, and the accuracy of the MNIST test data is captured at every 100 epochs. In the dDSC model, the data at each local site are sampled with a separate GMM process for each class, and the distributions are passed to every other site. The local sites then sample 20,000 images from each of the incoming distributions. We validate the procedure by testing each site’s model on the MNIST testing set and averaging the accuracies across every site.

In the second MNIST experiment for both dDS-NN and dDS-SVM, we bias the per-site data by class label. One site had access to digits 0-2, the second site had digits 4-6, and the last site contained digits 7-9. As the number three is not included, there are only 53,869 total images. As with the first experiment, each site using the distribution-sampled model sampled from the other sites’ per-class distributions of the respective local datasets. Again, the calculated accuracies per local site are averaged to encompass possible error differences between the sites.

The third experiment, which was only used to test the dDS-NN and not the dDS-SVM, uses 60,000 images as in the first experiment. The data is processed the same way as the first experiment, and the models are also of the same architecture. However, the primary difference is that the data is separated into 20 local sites as opposed to 3. This means that there are a total of 3,000 images at each local site distributed uniformly at random. Then, as in the previous experiments, the accuracies between the three models are compared.

The dDS-SVM model was tested on the same MNIST dataset as was used to test the dDS-NN. In the first SVM experiment, we randomly and uniformly distributed all of the training data across 3 sites. The second SVM experiment used 3 sites and the data was biased by label such that each site had exclusive access to 3 of the digits. Monte Carlo cross-validation was again used for both of these experiments.

### 2.6 sMRI data demographics

The sMRI dataset is aggregated from three separate datasets: the mind clinical imaging consortium (MCIC) [Gollub et al., 2013], the function biomedical informatics research network (fBIRN) [Potkin et al., 2008], and the center for biomedical research excellence (COBRE). In total, the three datasets consists of 584 subjects, of which, 269 are schizophrenia patients; the remaining subjects are healthy controls. For a more detailed breakdown of the demographic information, please see table 2.

**Table 2:** sMRI Subject Demographics

|  | MCIC | COBRE | FBIRN |
| --- | --- | --- | --- |
| Schizophrenia Subjects | 101 | 93 | 75 |
| Healthy Subjects | 119 | 97 | 99 |
| Schizophrenia Male/Female Split | 74/27 | 76/17 | 63/12 |
| Healthy Male/Female Split | 73/46 | 69/28 | 69/30 |
| Total | 220 | 190 | 174 |

### 2.7 sMRI dataset experiments

The sMRI dataset is very large with over 58 thousand features. As such, with or without statistical inference, the network traffic would be problematic. Due to this, we use a diagonal matrix to store the estimated variances of the GMM. Diagonal matrices store fewer data points and greatly reduces the required network traffic as it requires *N* values, whereas full covariance matrices require *N* ^2^ values. However, this loss of information may also have an impact on accuracy.

The sMRI dataset is analyzed by the dDS-NN using two separate experiments: a test in which the entire dataset is randomized across all local sites, and an experiment in which the local site data are biased by class. For this dataset, both experiments used four sites, but due to the size of the dataset, 10-fold cross-validation is used to measure the efficacy of the model. For all experiments of the neural network, we use two hidden layers of 1000 nodes and 200 nodes respectively. The batches are normalized and the weight sets are initialized with the Xavier algorithm, and we used a learning rate of 0.01. The models are run for a total of 2000 epochs throughout the entire dataset and the accuracy is measured on the per-fold test data every 100 epochs. In the randomized local dataset experiment, the three approaches operate the same way as in the MNIST experiment.

In the second dDS-NN experiment, we bias the per-site data by class label. The schizophrenia patients are evenly and randomly distributed between two of the local sites. The healthy controls are evenly and randomly split between the remaining two sites. No subject overlaps between the local sites, and each site had access to data with only one label; either schizophrenia or healthy. The distributions for each local dataset are then calculated and passed to the remaining local sites and each site trained a model on the sampled data and available local data. The local models are built with the same architecture, but since they are trained on different data they are not perfectly similar the way the local dDNN models are. This is measured by averaging the accuracies across all four local sites. The goal is to show the model’s robustness to extremely biased data.

We also use the sMRI data to test the dDS-SVM by uniformly distributing the sMRI data across four sites at random. 10-fold cross-validation was used to test the entire dataset. This was compared to an centralized SVM in which one site has access to the entire dataset.

## 3 Results

### 3.1 MNIST experiments

The first experiment of the neural network, in which the entire dataset is randomly distributed across all three sites, shows near identical accuracies between the dDNN and centralized approaches. Both converge at approximately 97.1% accuracy (Figure 2). The dDS-NN slightly worse, but still quite well, approximately converging towards 96.4% accuracy.

**Figure 1:**
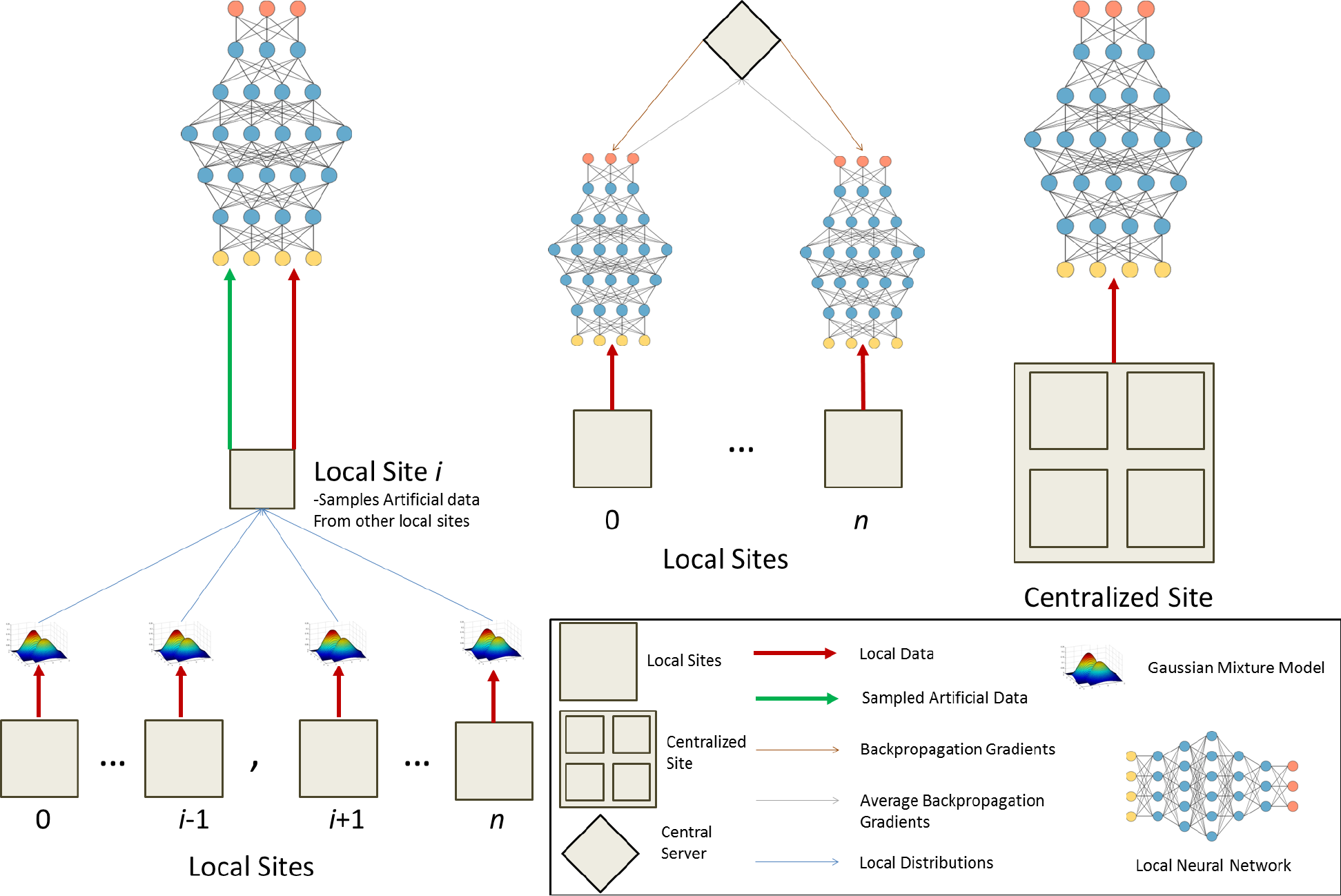
The three paradigms from left to right: dDS-NN, DDNN, and a centralized model. In the dDSC model, for every site *i*, every other site calculates the distribution of the local data and passes the distribution (in the form of a matrix) to site *i*. Site *i* then samples data from these distributions and uses this artificial data as well as the local data to train its own model. In the dDNN model, each local site trains its own model and the available local data and passes the gradient data to a centralized server. The centralized server then averages the local gradients and passes this average to the local sites to train the local models. The centralized paradigm uses all possible data in a single model at a central site.

**Figure 2:**
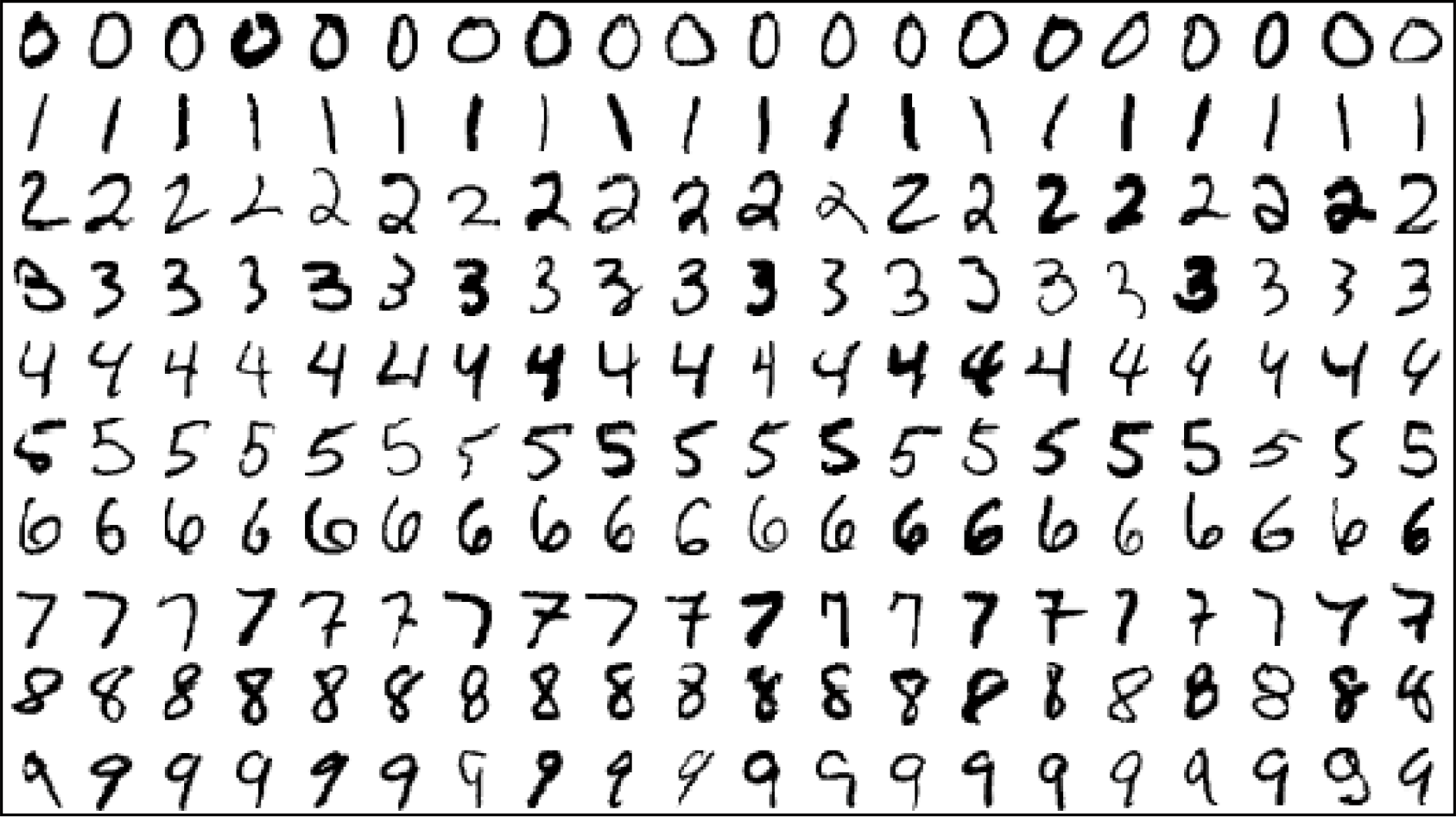
Examples of the hand-drawn digits in the MNIST dataset.

**Figure 3:**
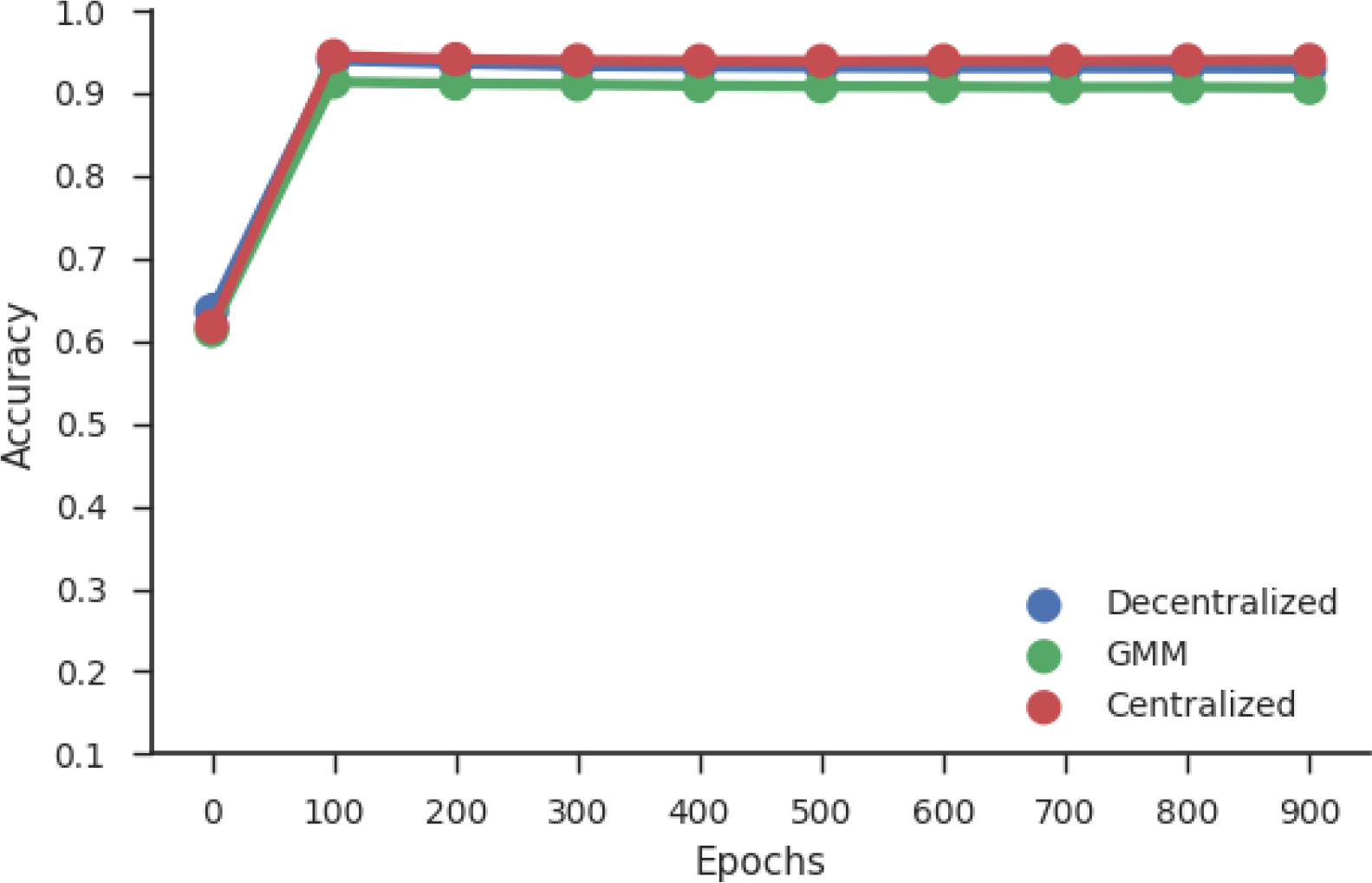
The MNIST data are randomly and evenly distributed between three sites for all three approaches, a centralized neural network, the dDNN model, and the dDS-NN model. The accuracy per approach is outlined above. The Centralized and dDNN methods are near identical.

In the second neural network experiment, the data are biased in such a way that each site had access to only three of the possible classes. Digit ‘3’ is removed so as to give each site an equal number of classes. From Figure 5, we see that, as in the previous experiment, the dDNN and centralized approaches are almost identical, converging toward 97.8%. The dDS-NN approach is slightly less accurate, converging towards 95.5% accuracy. The discrepancy between the accuracies of the same procedures is most likely due to the impact from the exclusion of digit three. It appears that including digit three makes the MNIST problem slightly more difficult.

**Figure 4:**
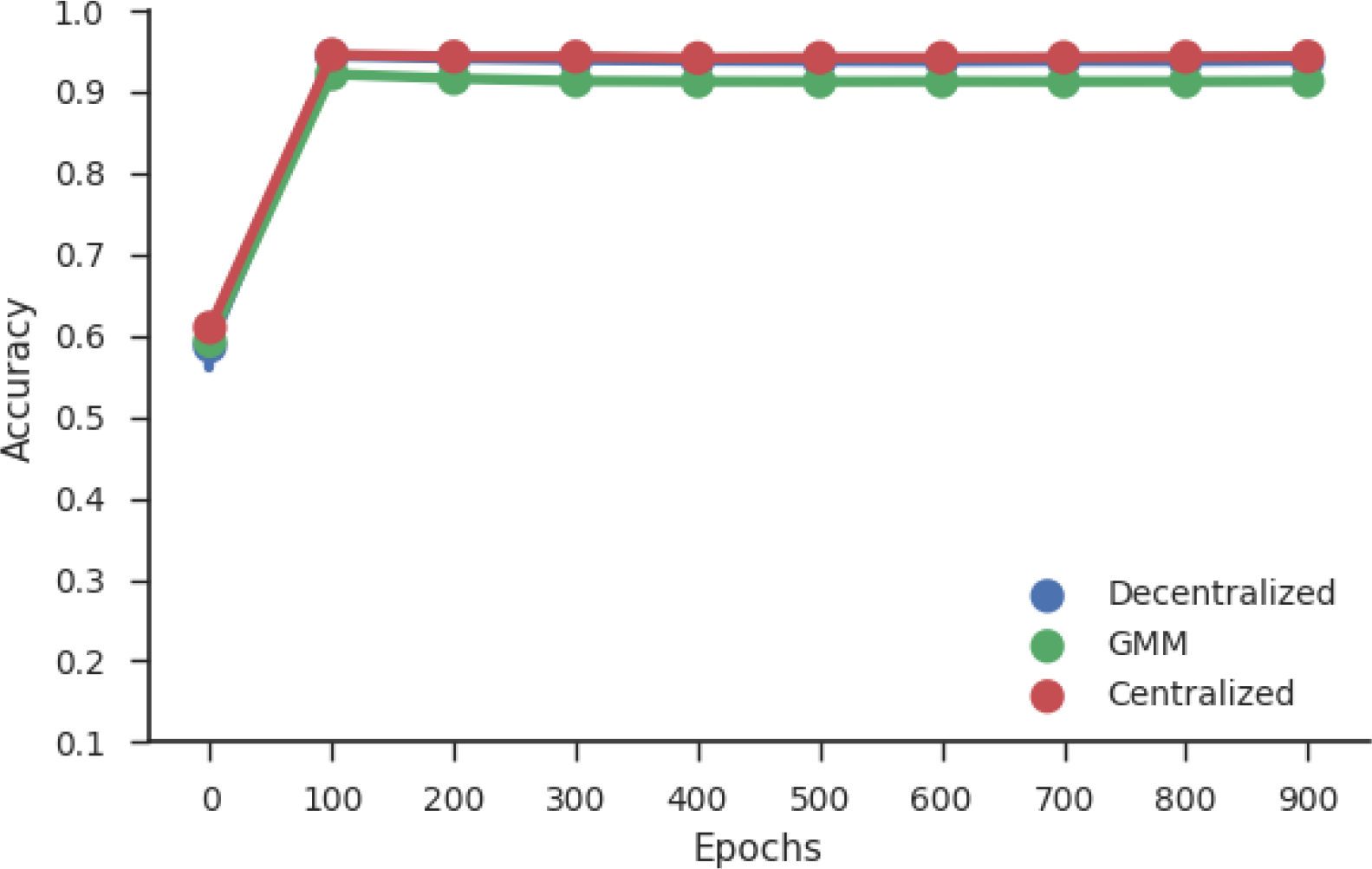
Results comparing the Centralized, dDNN, and dDSC approaches when the local MNIST data are biased by class label.

**Figure 5:**
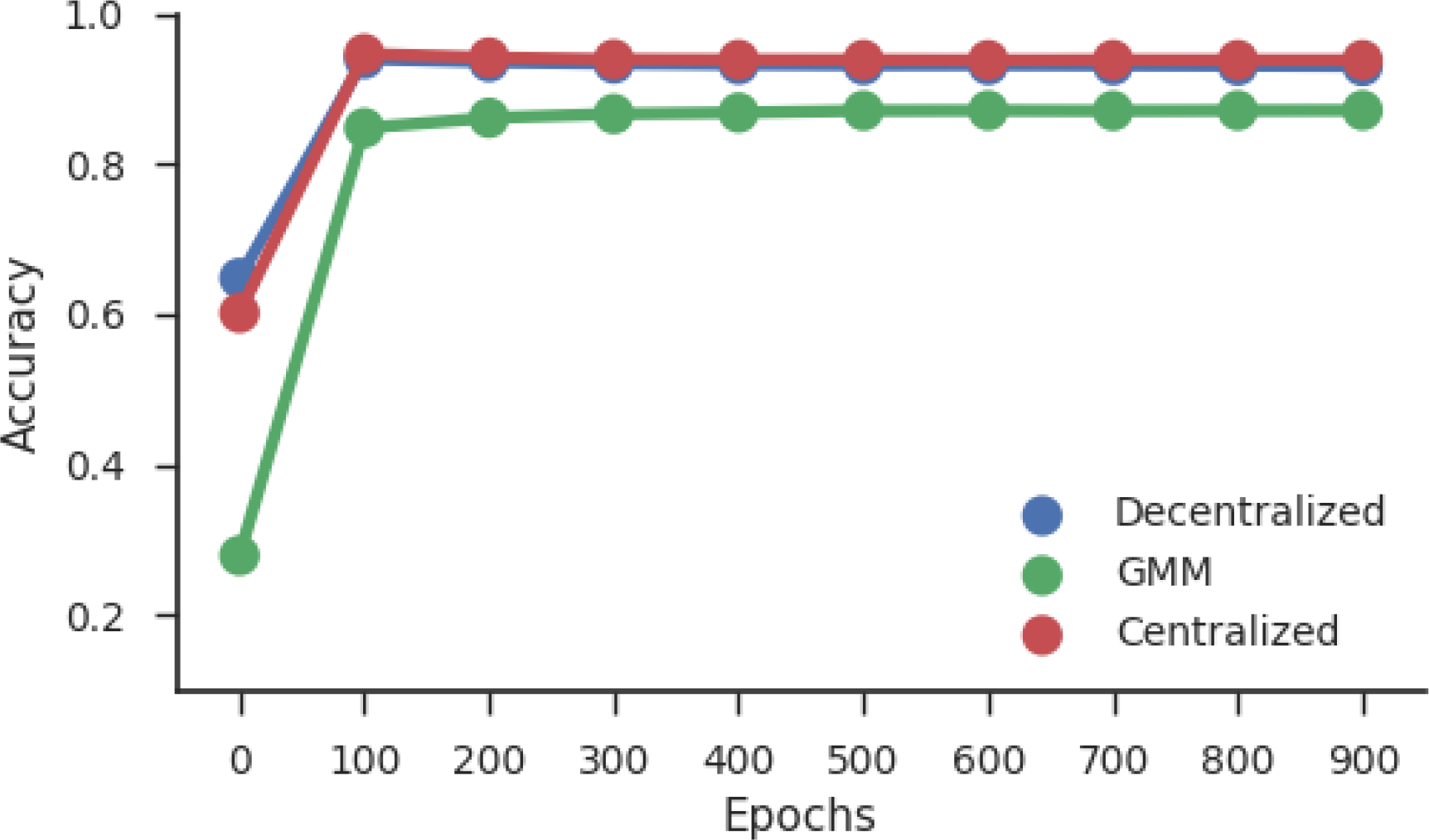
The MNIST data are randomly and evenly distributed between 20 sites for all three approaches. The accuracy per approach is outlined above.

The third neural network experiment shows the limitations of the dDS-NN model. This model converges towards 92%. Whereas the centralized and DDNN are almost identical and converges toward approximately 96.4% (Figure 2). The goal of this experiment is to show a major limitation in the procedure; which is its poor performance with many local sites.

The dDS-SVM performs as well as the dDS-NN compared to their centralized counter-parts in the case in which all of the data are distributed uniformly at random to all 3 sites. The average accuracy for the three site case for the dDS-SVM model is 91%. This is in contrast to the centralized method which had an accuracy of 91.5%. These accuracies reflect the differences between SVMs and neural networks when applied to the MNIST dataset [Deng, 2012].

For the case in which there are 3 sites and the data are biased by class label, the dDS-SVM had an accuracy of 90.8%, while the centralized method had an accuracy of 91.5%.

### 3.2 sMRI experiments

For all sMRI experiments, a diagonal covariance matrix is used to estimate the parameters of the distribution. This would have an impact on the performance of a model, which can be seen in the results. However, a diagonal covariance matrix greatly reduces the required network traffic from O(*N*^2^ * *L*) to O(*N* * *L*). Although the accuracy does decline by a small factor, the multishot dDNN and SVM models require a much higher order of network traffic.

The first dDS-NN experiment, in which the entire dataset is randomly distributed across all three sites, show near identical accuracies between the dDNN and centralized approaches. Both converge similarly: the dDNN method converges to about 72.1% accuracy (Figure 6), averaged across all 10 folds, and the centralized approach converges to about 72.9% accuracy. The dDS-NN converges toward 68.5% accuracy.

**Figure 6:**
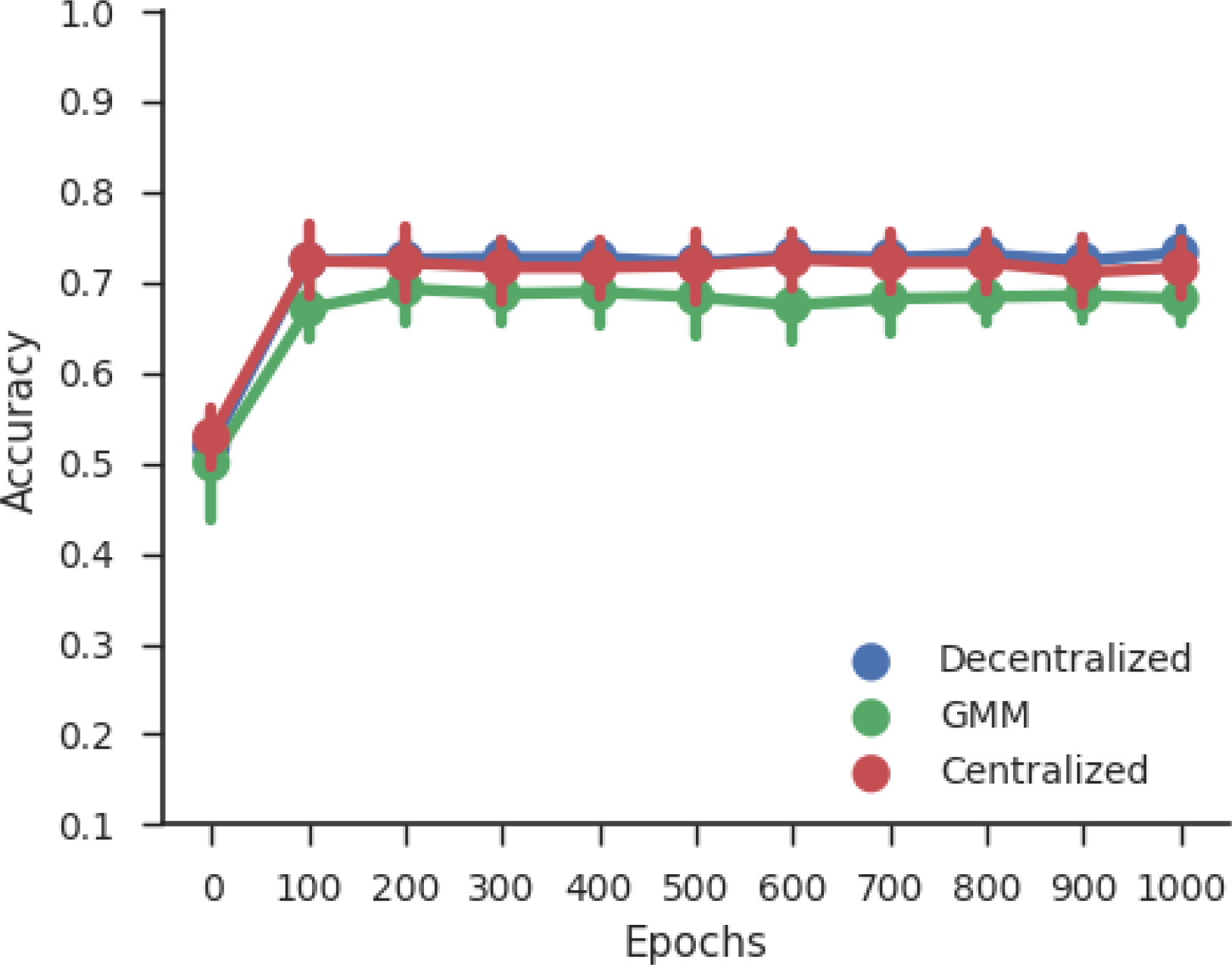
Results comparing the 10-fold cross-validation accuracies between Centralized, dDNN, and dDS-NN approaches with randomly and uniformly distributed data from the sMRI dataset. Stems show standard deviation and the lines show the mean accuracies across the 10 folds for each approach.

In the second sMRI experiment of the dDS-NN model, the data is evenly distributed across four sites, but is biased in such a way that each site had access to only one of the possible classes. This means that two sites had access to only patients and the remaining two sites had access to controls. From Figure 7, we see that, as in the previous experiment, the dDNN and centralized approaches are almost identical, converging towards 72.8% accuracy. The dDS-NN approach converges towards 65.1% accuracy.

**Figure 7:**
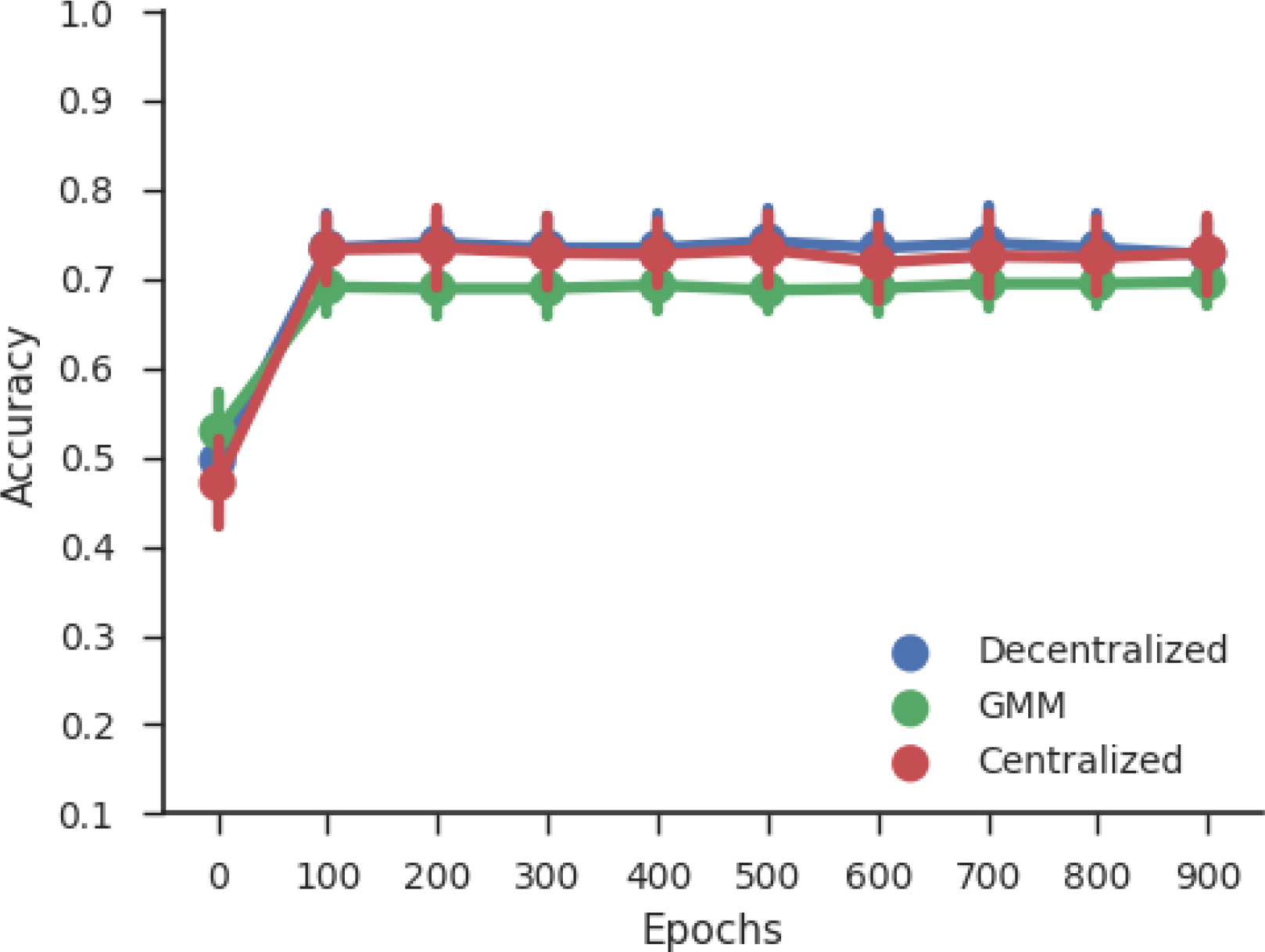
Results comparing the 10-fold cross-validation accuracies between Centralized, dDNN, and dDS-NN approaches with data from the sMRI dataset in which the data are biased by class label. Stems show standard deviation and the lines show the mean accuracies across the 10 folds for each of the three approaches.

**Figure 8:**
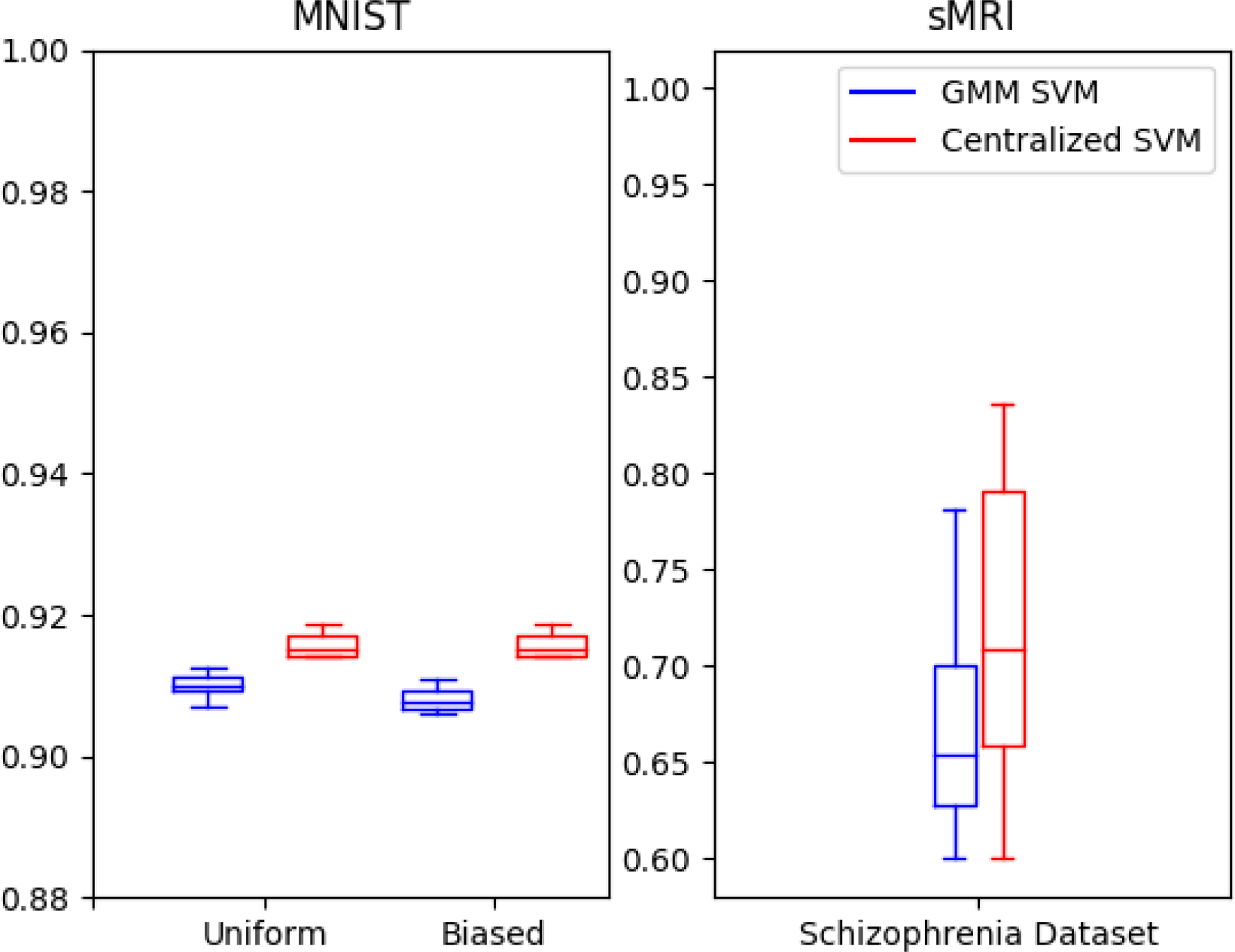
Results from the SVM experiments. The MNIST data (left) showed similar results between the centralized and dDS-SVM models. The uniformly and randomly distributed experiment of the MNIST dataset is on the far left and the case in which the data is biased by class label is in the center. The dDS-SVM also showed similar results compared to a centralized model when applied to the sMRI data (right).

When the sMRI data is modeled with an SVM, we uniformly and randomly distribute the data across 3 sites. 10-fold cross validation is used, covering all data samples, and the data are uniformly distributed to four different sites at random. The mean of the accuracies across all 10 folds from the dDS-SVM model is 67.5%, whereas the mean from the centralized method across all folds is 72%. These results are congruent with the results from the dDS-NN.

### 3.3 Network Traffic Analysis

The importance of this work rests in the dDSC model’s ability to reduce total network traffic. Our asymptotic analysis of the models provides a method to quantify the total network load as a function of the magnitude of the input data. Asymptotic analysis is a widely used method to understand the speed or space constraints of algorithms [de Bruijn, 1959]. The asymptotics are computed as the big-O, or the upper bound on the total amount of space required with respect to a given input of size *N*. This is done simply enough by computing the upper bound of how many gradients the multishot methods must pass between sites and how often this transfer must occur. The asymptotics of our singleshot method are estimated as a function of the size of the distributions passed between sites, including the means of each component and the covariance matrices of the local data.

The dDNN model proves to be very network intensive, as all gradients are passed between sites for every iteration. In order to reduce the total number of iterations required, the dDNN would train on one epoch ever iteration, which is analogous to stochastic gradient descent. Given this, the asymptotic analysis of the network traffic is straight forward: O(*W* * *I*), where *W* is the total number of weights in the model, and *I* is the number of iterations the model requires to converge. Given that the first weight matrix is size *N* \**w*_1_, *W* ≥ *N* for any possible *N*. With complex datasets, the number of weights can be much higher than N, as even first weight set, *w_i_*, is a multiplicative combination of the number of input features and the number of hidden nodes, and the number of hidden nodes can be on the order of size *N*, and the subsequent weight set is a multiplicative combination of *w*_1_. This suggests an upper bound of O(*N*^2^). It also must be noted that this network traffic occurs for every local site twice; the first time when the local sites pass gradient information to the central server, and the second time when the central server passes the averaged gradients to the local sites. This network load occurs every iteration *I* through the local dataset, where each iteration can be small batches of data, or an entire epoch of the local datasets. This means that the total space complexity of the multishot neural network is O(*N*^2^ * *I*). As neural networks can require many hundreds or even thousands of iterations, it grows as complexity of the data and model increases.

The current decentralized SVM model calculates the support vector weights, *v_i_* for every site *i* and broadcasts *v_i_* to all other local sites in a given iteration. Every feature *n* has a corresponding *v_i_* weight value, meaning *v_i_* is of size *N* for every *v* ∈ *V*. With *I* iterations and *N* input features, the space complexity of an SVM is O(*N* * *I* * *L*). The dDSC model however, requires statistical information to be passed only once. When a full covariance matrix is used to estimate the distribution of the GMM, the model requires space for the variance of each feature pair, for each possible class. Meaning, at most, O(*N* ^2^ * *L*) bytes of information are passed to each other site for each site with a number of labels L. However, in the case of a diagonal matrix, which is the method used to analyze the sMRI data, the variances and means of only the individual features are stored in memory for each label, meaning this method requires O(*N* * *L*) traffic across a network.

Given this analysis, it is clear why the dDSC model is effective at reducing the upper bound of the total network traffic, as well as greatly reducing the number of times information is passed between sites. Our analysis showed that our singleshot method does decrease network traffic by at least one order of magnitude. Beyond the reduction in total network traffic, it is important to note that the dDSC model reduces the total number of network broadcasts required during training. With multiple network broadcasts, there are more fault points, as any interruption in the network connection halts training, and the probability of interruption increases as the number of required broadcasts increases. This problem is entirely mitigated with a singleshot method.

## 4 Discussion

Decentralized models have an important place in machine learning and data analytics and are based on the concept of distributed computation [Dijkstra, 1965, Dean et al., 2012]. Currently, decentralized modeling has focused on basic machine learning techniques [Gazula et al., 2018, Ming et al., 2017, Plis et al., 2016, Baker et al., 2015, Forero et al., 2010], with scant research on neural networks [Lewis et al., 2017]. Previous research on decentralized deep learning focused on eliminating significant differences in accuracy between decentralized neural networks and their centralized counterpart. However, this proved to be problematic as the size of the network traffic is unmanageable for even the fastest network, given a large enough dataset, such as fMRI scans for many subjects. As the research is geared towards biological data, consisting of very large datasets, this problem is especially important.

Mixture models have previously been used to enhance neural networks [Viroli and McLachlan, 2017], and the concept of generating new samples to improve neural network accuracy has also been explored [He et al., 2008, Goodfellow et al., 2014]. We adapt these concepts to greatly improve decentralized neural networks and solve the problem of high network traffic with minimal impact on accuracy.

Overall, the dDSC model drastically decreased network flow by a polynomial amount. The multishot neural network’s asymptotic space requirement is O(*W* * *I*) for every site, where *W* has a magnitude of greater than or equal to the size of the input, *N*. While the multishot SVM requires O(*N* * *I* * *L*) network load. This contrasts with our singleshot implementation which requires only O(*N*) space.

Although the model appears to be valid for many criteria, it does have limitations. As seen in Figure 2, the accuracy of the dDSC model decreases with many more local sites. The local distributions appear to be less accurate due to a lack of sample data. This result is what would be expected for small enough sample sizes. However, there is a body of literature showing how to improve mixture model accuracy given small datasets [Liu et al., 2008]. Although, it should be noted that small datasets will always be a limiting factor for machine learning, and a certain amount of error is expected with small datasets.

We suggest that future researchers could develop models that are more accurate in general, or at least in the case of many local sites. This could include testing different mixture models or methods to generate artificial data to improve the accuracy of local distributions for small datasets [Fei-Fei et al., 2007]. Future research could also focus on using generative adversarial networks (GAN) to train on local data and produce artificial samples for other local sites. Finally, a more basic avenue for continued research would be to use our method for different neural network architectures or different machine learning models. This model shows that statistical inference works as a way to decentralize data for classifiers. However, statistical inference could also be used for other decentralized methods.

## 5 Conclusion

This work expands on the current body of decentralized classification models with a technique to improve network efficiency by estimating distributions of the local data and sampling from these distributions at every other local site. We accomplished this with a Gaussian mixture model to produce artificial samples from the local data. The results from the experiments show a promising expansion to decentralized neural networks. We showed that the dDS-NN produces accuracy comparable to the dDNN model, but with greatly reduced network traffic. This research also concludes that the dDS-SVM performs similarly to a centralized SVM while also reducing network traffic compared to the original decentralized SVM when using a diagonalized matrix.

## References

Aine, C. J., Bockholt, H. J., Bustillo, J. R., Cañive, J. M., Caprihan, A., Gasparovic, C., Hanlon, F. M., Houck, J. M., Jung, R. E., Lauriello, J., Liu, J., Mayer, A. R., Perrone-Bizzozero, N. I., Posse, S., Stephen, J. M., Turner, J. A., Clark, V. P., and Calhoun, V. D. (2017). Multimodal neuroimaging in schizophrenia: Description and dissemination. Neuroinformatics, 15(4):343–364.

Baker, B. T., Silva, R. F., Calhoun, V. D., Sarwate, A. D., and Plis, S. M. (2015). Large scale collaboration with autonomy: Decentralized data ica. In 2015 IEEE 25th International Workshop on Machine Learning for Signal Processing (MLSP).

Carter, K. W., Francis, R. W., Carter, K., Francis, R., Bresnahan, M., Gissler, M., Grønborg, T., Gross, R., Gunnes, N., Hammond, G., et al. (2015). Vipar: a software platform for the virtual pooling and analysis of research data. International journal of epidemiology, 45(2):408–416.

de Bruijn, N. G. (1959). Asymptotic methods in analysis. Bull. Amer. Math. Soc, 65(3):160–163.

Dean, J., Corrado, G., Monga, R., Chen, K., Devin, M., Mao, M., aurelio Ranzato, M., Senior, A., Tucker, P., Yang, K., Le, Q. V., and Ng, A. Y. (2012). Large scale distributed deep networks. In Pereira, F., Burges, C. J. C., Bottou, L., and Weinberger, K. Q., editors, Advances in Neural Information Processing Systems 25, pages 1223–1231. Curran Associates, Inc.

Deng, L. (2012). The mnist database of handwritten digit images for machine learning research. IEEE Signal Processing Magazine, 29(6):141–142.

Dijkstra, E. (1965). Solution of a problem in concurrent programming control. Communications of the ACM, 8:569.

Fei-Fei, L., Fergus, R., and Perona, P. (2007). Learning generative visual models from few training examples: An incremental bayesian approach tested on 101 object categories. Computer Vision and Image Understanding, 106(1):59–70. Special issue on Generative Model Based Vision.

Forero, P. A., Cano, A., and Giannakis, G. B. (2010). Consensus-based distributed support vector machines. J. Mach. Learn. Res., 11:1663–1707.

Gazula, H., Baker, B. T., Damaraju, E., Plis, S. M., Panta, S. R., Silva, F., and Calhoun, V. D. (2018). Decentralized analysis of brain imaging data: Voxel-based morphometry and dynamic functional network connectivity. Frontiers in neuroinformatics, 12.

Gollub, R. L., Shoemaker, J. M., King, M. D., White, T., Ehrlich, S., Sponheim, S. R., Clark, V. P., Turner, J. A., Mueller, B. A., Magnotta, V., et al. (2013). The mcic collection: a shared repository of multi-modal, multi-site brain image data from a clinical investigation of schizophrenia. Neuroinformatics, 11(3):367–388.

Goodfellow, I., Pouget-Abadie, J., Mirza, M., Xu, B., Warde-Farley, D., Ozair, S., Courville, A., and Bengio, Y. (2014). Generative adversarial nets. In Ghahramani, Z., Welling, M., Cortes, C., Lawrence, N. D., and Weinberger, K. Q., editors, Advances in Neural Information Processing Systems 27, pages 2672–2680. Curran Associates, Inc.

Hanlon, F. M., Houck, J. M., Pyeatt, C. J., Lundy, S. L., Euler, M. J., Weisend, M. P., Thoma, R. J., Bustillo, J. R., Miller, G. A., and Tesche, C. D. (2011). Bilateral hippocampal dysfunction in schizophrenia. Neuroimage, 58(4):1158–1168.

He, H., Bai, Y., Garcia, E. A., and Li, S. (2008). Adasyn: Adaptive synthetic sampling approach for imbalanced learning. In 2008 IEEE International Joint Conference on Neural Networks (IEEE World Congress on Computational Intelligence), pages 1322–1328.

LeCun, Y., Cortes, C., and Burges, C. (2010). Mnist handwritten digit database. AT&T Labs [Online]. Available: http://yann.lecun.com/exdb/mnist, 2.

Lewis, N., Plis, S., and Calhoun, V. (2017). Cooperative learning: Decentralized data neural network. In Neural Networks (IJCNN), 2017 International Joint Conference on, pages 324–331. IEEE.

Liu, Y., Hayes, D. N., Nobel, A., and Marron, J. S. (2008). Statistical significance of clustering for high-dimension, low–sample size data. Journal of the American Statistical Association, 103(483):1281–1293.

Ming, J., Verner, E., Sarwate, A., Kelly, R., Reed, C., Kahleck, T., Silva, R., Panta, S., Turner, J., Plis, S., et al. (2017). Coinstac: Decentralizing the future of brain imaging analysis. F1000Research, 6.

Plis, S. M., Sarwate, A. D., Wood, D., Dieringer, C., Landis, D., Reed, C., Panta, R., Turner, J. A., Shoemaker, J. M., Carter, K. W., et al. (2016). Coinstac: a privacy enabled model and prototype for leveraging and processing decentralized brain imaging data. Frontiers in neuroscience, 10:365.

Potkin, S., Turner, J., Brown, G., McCarthy, G., Greve, D., Glover, G., Manoach, D., Belger, A., Diaz, M., Wible, C., et al. (2008). Working memory and dlpfc inefficiency in schizophrenia: the fbirn study. Schizophrenia bulletin, 35(1):19–31.

Saha, D. K., Calhoun, V. D., Panta, S. R., and Plis, S. M. (2017). See without looking: joint visualization of sensitive multi-site datasets. In Proceedings of the Twenty-Sixth International Joint Conference on Artificial Intelligence (IJCAI’2017), pages 2672–2678.

Thompson, P. M., Stein, J. L., Medland, S. E., Hibar, D. P., Vasquez, A. A., Renteria, M. E., Toro, R., Jahanshad, N., Schumann, G., Franke, B., et al. (2014). The enigma consortium: large-scale collaborative analyses of neuroimaging and genetic data. Brain imaging and behavior, 8(2):153–182.

Viroli, C. and McLachlan, G. J. (2017). Deep gaussian mixture models. CoRR, abs/1711.06929.

Wojtalewicz, N. P., Silva, R. F., Calhoun, V. D., Sarwate, A. D., and Plis, S. M. (2017). Decentralized independent vector analysis. In Acoustics, Speech and Signal Processing (ICASSP), 2017 IEEE International Conference on, pages 826–830. IEEE.

